# Accumulated metagenomic studies reveal recent migration, whole genome evolution, and undiscovered diversity of orthomyxoviruses

**DOI:** 10.1101/2022.08.31.505987

**Authors:** Gytis Dudas, Joshua Batson

## Abstract

Metagenomic studies have uncovered many novel viruses by looking beyond hosts of public health or economic interest. However, the resulting viral genomes are often incomplete, and analyses largely characterize the distribution of viruses over their dynamics. Here, we integrate accumulated data from metagenomic studies to reveal geographic and evolutionary dynamics in a case study of *Orthomyxoviridae*, the RNA virus family that includes influenza virus. First, we use sequences of the orthomyxovirid Wǔhàn mosquito virus 6 to track the migrations of its host. We then look at orthomyxovirus genome evolution, finding gene gain and loss across members of the family, especially in the surface proteins responsible for cell and host tropism. We find that the surface protein of Wǔhàn mosquito virus 6 exhibits accelerated non-synonymous evolution suggestive of antigenic evolution *i*.*e*. vertebrate infection, and belongs to a wider quaranjavirid group bearing highly diverged surface proteins. Finally we quantify the progress of orthomyxovirus discovery and forecast that many diverged *Orthomyxoviridae* members remain to be found. We argue that continued metagenomic studies will be fruitful for understanding the dynamics, evolution, ecology of viruses and their hosts, regardless of whether novel viruses are identified or not, as long as study designs allowing for the resolution of complete viral genomes are employed.

**Importance:** The number of known virus species has increased dramatically through metagenomic studies, which search genetic material sampled from a host for non-host genes. Here, we focus on an important viral family that includes influenza viruses, the *Orthomyxoviridae*, with over a hundred recently discovered viruses infecting hosts from humans to fish. We find one virus called Wǔhàn mosquito virus 6, discovered in mosquitoes in China, has spread across the globe very recently. Surface proteins used to enter cells show signs of rapid evolution in Wǔhàn mosquito virus 6 and its relatives which suggests an ability to infect vertebrate animals. We compute the rate at which new orthomyxovirus species discovered add evolutionary history to the tree of life, predict that many viruses remain to be discovered, and discuss what appropriately designed future studies can teach us about how diseases cross between continents and species.

## 1 Introduction

### 1.1 Metagenomic virus discovery efforts to date

Viruses that cause disease in humans and economically important organisms were the first to be isolated and characterized. Recently, cheap DNA sequencing has enabled a wave of metagenomic studies in a broader range of hosts, in which viruses are identified in a host sample by nucleic acid sequence alone and a new virus is said to be discovered if that sequence is sufficiently diverged. As a result, the number of known viruses has increased by more than an order of magnitude in the decade since 2012 (Roux et al., 2021). While some entirely new viral families have been proposed (Obbard et al., 2020), many of these new viruses are interleaved on the virus tree of life with viruses infecting hosts of economic importance. Studying their ecology (Shi et al., 2019) and host associations (Li et al., 2015; Shi et al., 2018) provides insight into the host-switching and genome evolution processes important for the evolution of pathogenicity.

This richer tree of viruses has provided some early success stories, such as jingmenvirids that are currently associated with human disease (Wang et al., 2019) but which were first discovered metagenomically in ticks (Qin et al., 2014) indicating a likely route of transmission. Surveillance in hosts known to pose disproportionate risk, such as bats, (Ge et al., 2016) has similarly provided context for zoonotic pathogens like SARS-CoV-2 (Wu et al., 2020). Metagenomic studies carried out at scale can effectively multiplex other tasks previously addressed with targeted sampling, like understanding the evolutionary history of human pathogens (Keele et al., 2006) or using viruses that evolve faster than their hosts to track host movements (Wheeler et al., 2010).

### 1.2 Going beyond mere virus discovery

Here, we seek to show how accumulated data from metagenomic studies can provide deep insights into viral evolution and dispersion across a moderately diverged viral clade through a case study of *Orthomyxoviridae. Orthomyxoviridae* is a family of enveloped segmented negative sense single-stranded RNA viruses that infect vertebrates and arthro-pods. Traditionally it is split into genera *Thogotovirus, Quaranjavirus* (both mostly tick-borne and occasionally infecting vertebrates), *Isavirus* (found only in fish so far), and a genus for each of influenza A, B, C, and D viruses (found in various vertebrates, in-cluding humans). Orthomyxovirus discovery has historically been driven by impact on human health (*e*.*g*. influenza virus) and livelihood (*e*.*g*. salmon infectious anemia virus), or association with known disease vectors (*e*.*g*. the tick-borne Johnston Atoll quaranja-and Thogoto thogotovirus). The metagenomic revolution has resulted in ten times more orthomyxovirids being discovered over the last decade than in the previous 79 years since the first orthomyxovirus discovery, of influenza A virus, in 1933. The vast majority of known orthomyxoviruses use one of two surface protein classes, with vertebrate-infectingonly members (influenza and isavirids) using one or more class I membrane fusion proteins derived from hemagglutinin-esterase-fusion (HEF) (Parry et al., 2020), sometimes delegating the receptor cleavage function upon exit from the host cell to a separate protein neuraminidase (NA). Meanwhile, arthropod-infecting orthomyxovirids (quaranja- and thogotovirids, which sometimes spill over into vertebrates) use a class III membrane fusion protein called gp64 (Garry and Garry, 2008). Orthomyxovirus genomes are known to have 6-8 segments, but many metagenomically discovered viruses in this group have incomplete genomes. To our knowledge, an inventory of surface protein class use and segment content of *Orthomyxoviridae* members is not yet available.

We start by showing how closely related virus sequences observed across numerous studies can reveal host spatial dynamics and virus microevolution, using the orthomyx-ovirid Wǔhàn mosquito virus 6 (WuMV-6). WuMV-6 was discovered in China in 2013 from a single PB1 sequence (Li et al., 2015). WuMV-6 belongs to a diverse clade of arthropod orthomyxovirids somewhat distantly related to members of the genus *Quaranjavirus*. WuMV-6 has since turned up in *Culex* mosquitoes across numerous metagenomic studies all around the world (Shi et al., 2017; Pettersson et al., 2019; Batson et al., 2021). The abundance of WuMV-6 genome data and its amenability to molecular clock analyses are the main reasons why we chose to focus on it. To our knowledge WuMV-6 has not been isolated. Looking beyond WuMV-6, we map out known genome composition across members of *Orthomyxoviridae*, highlighting parts of the tree where changes to segment numbers are likely to have taken place. In this task we focus on the PB1 protein of the heterotrimeric orthomyxovirid RNA-directed RNA polymerase (RdRp) complex for most of our analyses, since PB1 encodes the RdRp motif responsible for replicase activity (the two other proteins of the RdRp heterotrimer are PB2 and PA). In looking at genome composition we pay close attention to surface protein use across the PB1 tree, and focus particularly on gp64 proteins used by thogoto- and quaranjaviruses. We find surface proteins to be quite mobile across members of *Orthomyxoviridae* over evolutionary timescales and identify a clade of quaranjaviruses known to have acquired new segments using distinctly diverged gp64 proteins. Finally we borrow methods from macroevolutionary research to quantitatively assess the pace at which orthomyxovirus evolutionary history is being uncovered, finding that despite their already transformative effect, metagenomic discovery efforts are likely to continue to find substantially diverged members of *Orthomyxoviridae* for some time.

## 2 Results

### 2.1 WuMV-6 exhibits rapid population dynamics

Wǔhàn mosquito virus 6 (WuMV-6), a mosquito orthomyxovirus seen frequently across much of the world (Pettersson et al., 2019; Li et al., 2015; Shi et al., 2017) belongs to a clade related to the genus *Quaranjavirus* and has two extra segments compared to other quaranjavirids (Batson et al., 2021). WuMV-6 is also distinct from many other arthropod RNA viruses in being found very often yet exhibiting limited genetic diversity and a strong molecular clock signal (Supp. Figs. 1, 2, and 3), allowing the use of phylogenetic methods like the reassortment network (Müller et al., 2020). Fig. 1A (also Supp. Fig. 4) shows a reassortment network analysis of currently available complete WuMV-6 genomes (seven from Australia, 13 from California, three from Cambodia, one from China, three from Sweden) collected between 2006 and 2020, depicting relationships between segments, their reassortments with respect to each other and timings of both. We find that all WuMV-6 segments share a common ancestor within the last 60-odd years, which is not unusual for insect viruses (Webster et al., 2015) (Fig. 1A and Supp. Figs. 5 and 6), and that a more recent, potentially global, sweep is underway, with segments from six continents sharing a common ancestor in the last 20 years (Fig. 1B).

**Figure 1.**
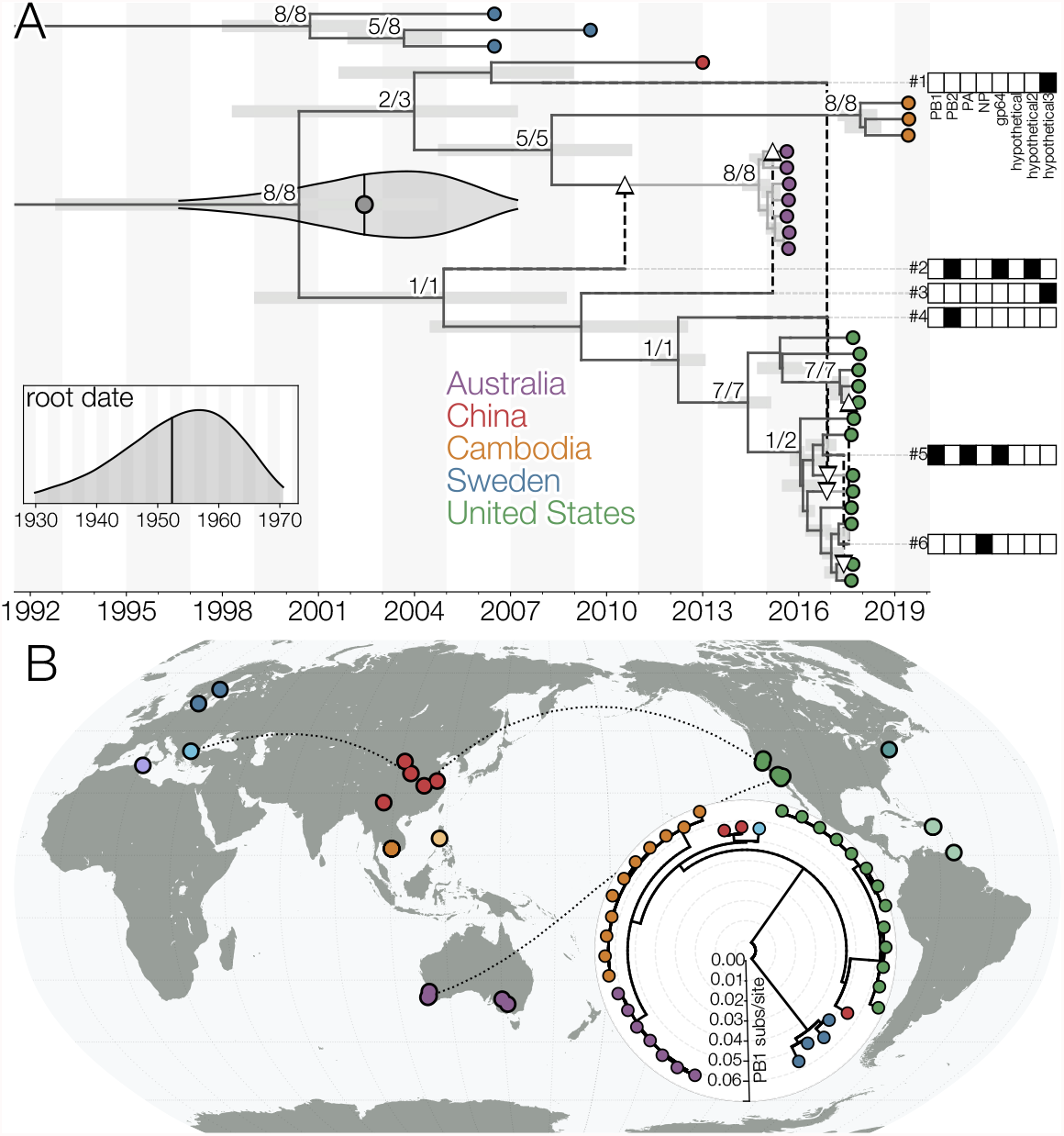
A) Reassortment network of complete WuMV-6 genomes. Tips are indicated with circles and colored based on location. Reassortant edges are indicated with dashed lines terminating in white arrowheads and numbered with segments carried along the edge indicated with filled-in rectangles to the right of the plot (for clearer segment embeddings see Supp. Fig. 4). The network is truncated to 1995 with a violin plot indicating the 95% highest posterior density (HPD) interval for the date of the common ancestor of non-Swedish WuMV-6 samples estimated from the posterior distribution of reassortment networks. Horizontal gray bars indicate node height 95% HPD intervals in the summary network. Black vertical line with the gray dot within the violin plot indicates the mean estimate. The inset plot indicates same for the root date of the network with black vertical line indicating the mean. Since the summary procedure for the posterior distribution of networks is overly conservative (see Supp. Fig. 5), node supports are expressed as number of times a given node is seen with ≥0.95 probability in segment embedding summary trees, after carrying out the subtree prune-regraft procedures for any given embedding indicated by reassortant edges, out of all such nodes. B) A maximum likelihood (ML) tree of WuMV-6 PB1 sequences, showing additional samples for which only incomplete genomes are available (Greece and China). Dots on the map indicate all locations where the presence of WuMV-6 has been detected, including complete (Australia, Cambodia, China, Sweden, USA) and incomplete genomes (China, and Greece), and detections at the read level (Connecticut, Philippines, Puerto Rico, Trinidad and Tobago, and Tunisia). Dotted lines connect locations that have experienced recent WuMV-6 gene flow based on reassortment patterns. For all available WuMV-6 segment data see Supp. Fig. 6.

Although the geographic population structure of WuMV-6 is appreciable, with samples from the same country often close on the tree, reassortment events indicate contact between genomic lineages across vast distances. For example, reassortment events #2 and #3 in Fig. 1A indicate contact as recently as 2010-2015 between WuMV-6 lineages eventually found in Australia and California. Similarly, some lineages found in China are related to recent (*circa* 2017) Californian lineages (reassortment event #1 in Fig. 1A). Even lineages not represented in the reassortment network due to incomplete genomes show evidence of gene flow, like Chinese and Greek PB1 sequences in Fig. 1B. These results indicate that WuMV-6 populations are very mobile.

### 2.2 Surface protein gp64 of WuMV-6 and its relatives evolves rapidly

By estimating the rates of synonymous and non-synonymous evolution we find that gp64, the surface protein of WuMV-6, is evolving faster in terms of non-synonymous substitutions per codon per year than the rest of the known WuMV-6 proteome, save for the smallest segment, which is expected to be spliced (hypothetical3 (Batson et al., 2021)) and therefore likely to contain overlapping reading frames (Fig. 2A and Supp. Fig. 7). The rate of non-synonymous evolution in WuMV-6 gp64 is also faster than the spike protein of endemic human coronaviruses (Kistler and Bedford, 2021) and about as fast as Ebola virus glycoprotein GP during the West African epidemic (Park et al., 2015)(Fig. 2A), with highest dN/dS values concentrated around its fusion loops (Garry and Garry, 2008) (Supp. Fig. 8). We see elevated rates of amino acid evolution in gp64 across the wider clade defined by Astopletus and Ū sinis viruses, to which WuMV-6 belongs. Members have PB1 proteins (encoding RdRp) closely related (Fig. 2B) but gp64 proteins substantially diverged from other quaranjaviruses and each other (Fig. 2C and Supp. Fig. 9). The pronounced non-synonymous divergence in gp64 at the WuMV-6 population level and the wider Asto-Ū sinis clade level indicates some evolutionary pressure on this surface protein, such as diversifying selection pressure from repeat infections of hosts with humoral immune systems.

**Figure 2.**
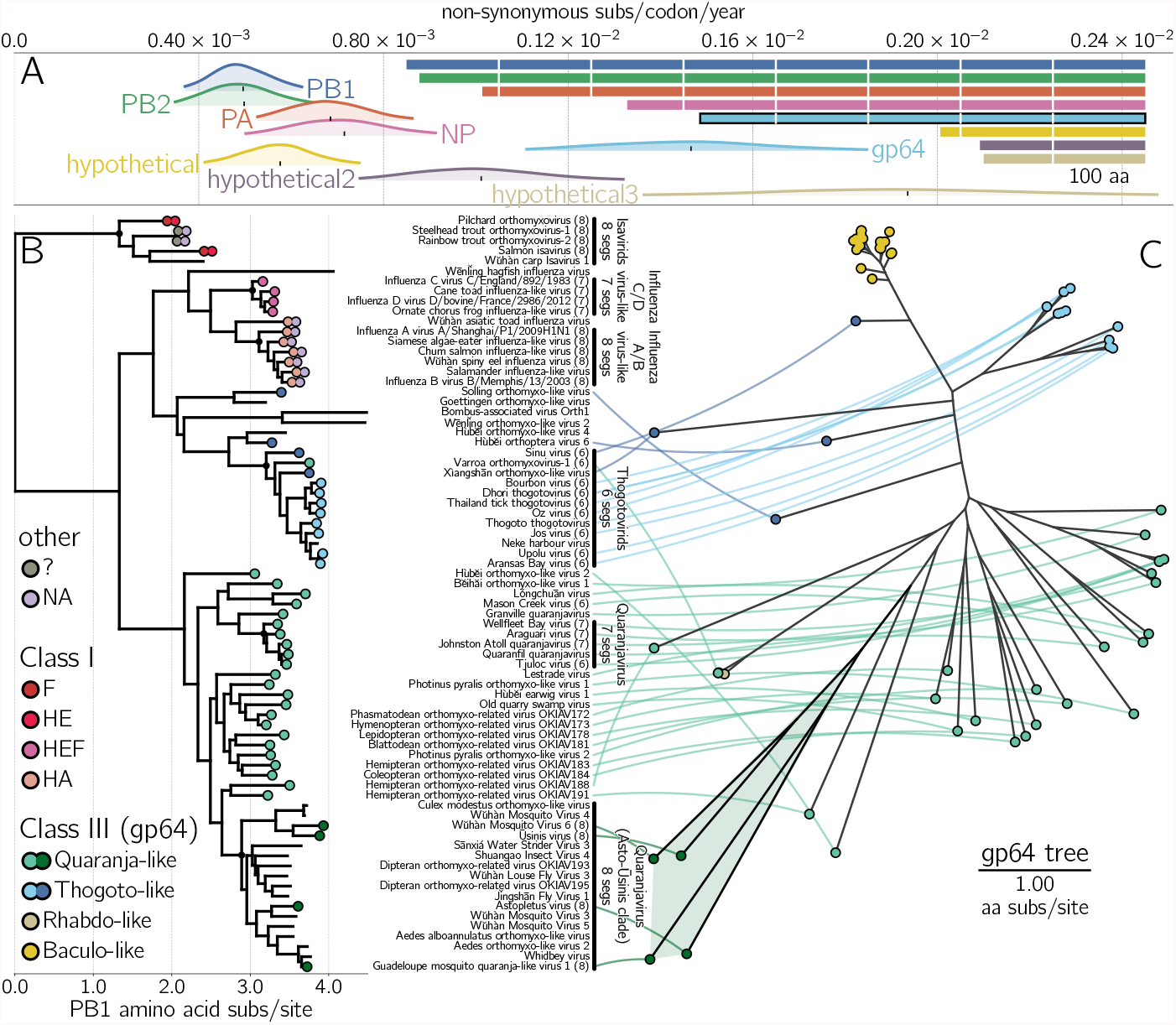
A) 95% highest posterior densities for the rate of non-synonymous mutations per codon per year for each known WuMV-6 gene (indicated by color). Black vertical ticks indicate the mean estimate. Lengths of open reading frames is shown to the right, with white lines denoting 100 amino acid increments and gp64 outlined in black. Note that putative hypothetical 3 protein is expected to be spliced (Batson et al., 2021) and may contain overlapping coding regions where synonymous changes in one frame may be non-synonymous in another. B) Rooted phylogenetic tree depicts the relationships between PB1 proteins of orthomyxoviruses. Surface proteins are marked as colored circles at the tips: red hues for class I membrane fusion proteins, green/blue and yellow/brown for class III proteins (gp64) of *Orthomyxoviridae* and non-*Orthomyxoviridae*, respectively, and lilac for neuraminidases. Likely genome composition for certain groups are highlighted with black vertical lines to the right of the tree, with corresponding implied common ancestor with such organization marked with a black circle in the tree. C) An unrooted phylogeny to the right of the PB1 tree shows the relationships between gp64 proteins found in thogoto- or thogoto-like-(blue), quaranja- or quaranja-like-(green), baculo-(yellow), and some rhabdovirids (tan). Where available, each orthomyxovirid gp64 protein is connected to its corresponding PB1 sequence. Black branches with a faded green area in the gp64 tree to the right indicate the position of Asto-Ū sinis gp64 proteins.

Comparing the numbers of PB1 and gp64 proteins sequences discovered so far indicates a clear paucity of the latter, especially in the Asto-Ū sinis clade, which highlights the poor general state of knowledge of genome composition across members of *Orthomyxoviridae*. The closest relative with reliable segment information (based on EM photographs (Allison et al., 2015)) is the genus *Quaranjavirus* with seven segments (Fig. 2B), indicating that since the common ancestor of the *Quaranjavirus* genus and the Asto-Ū sinis clade, segments were either lost in the former or gained in the latter. Such gaps (Fig. 2B) in understanding seem to increase with phylogenetic distance from vertebrate-pathogenic viruses, a hallmark of retrospective research into outbreaks rather than prospective efforts aimed at understanding the correlates of pathogenicity across members of this family.

### 2.3 Surface proteins of *Orthomyxoviridae* are prone to horizontal gene transfer

The phylogenetic tree of PB1 with surface protein classes indicated also demonstrates the plasticity in orthomyxovirus genome composition — regardless of rooting, the PB1 tree requires at least two switches in viral membrane fusion protein class to explain the current distribution of HEF-like (class I) and gp64-like (class III) proteins. Even within gp64-using orthomyxoviruses changes between different gp64 lineages are apparent, *e*.*g*. Hùběi orthomyxo-like virus 2 carries gp64 related to the Asto-Ū sinis clade yet does not belong to it in PB1 (Fig. 2B and Supp. Fig. 9). Almost all orthomyxoviruses use either HEF-like or gp64-like proteins, with Rainbow / Steelhead trout orthomyxoviruses (and one additional relative (Batts et al., 2017), not shown) being the only exceptions. Both clearly possesses an influenza A/B virus-like neuraminidase (NA) but the protein termed “hemagglutinin” (Batts et al., 2017) does not resemble any known protein (Finn et al., 2011). While *Orthomyxoviridae* are a moderately-sized virus family, their members make use of a diverse and evolving set of surface proteins.

### 2.4 Rate of *Orthomyxoviridae* PB1 diversity discovery remains high

We now analyze the progress made by virus discovery studies on *Orthomyxoviridae* members by quantifying the evolutionary contribution (branch length in amino acid substitutions per site) contributed by each new orthomyxovirid taxon. There are two clear phases of discovery: before 2015, public health investigations of pathogens infecting humans and farmed animals or vectored by ticks led to the discovery of 14 viruses. Since 2015, when lower costs of sequencing enabled large metagenomic surveys in arthopods and vertebrates of little immediate economic value (Li et al., 2015; Shi et al., 2018), 115 additional viruses have been discovered (Supp. Fig. 10A) with current trends, when focusing on smaller clades, being consistent with arthropod discovery efforts being the most fruitful (Supp. Fig. 10B).

To quantify how each discovery contributed to our knowledge of the family’s evolutionary history, we take a phylogenetic approach, building a maximum-likelihood tree of the sole protein shared by all RNA viruses, RNA-directed RNA polymerase (RdRp) (Koonin and Dolja, 2014), encoded here in the PB1 gene (Kobayashi et al., 1996) (Fig. 3A). We scan through the tree based on the chronology of discovery, attributing to each taxon the sum of the lengths of the branches ancestral to that taxon but not to earlier taxa. This quantity is called the phylogenetic diversity (PD), a metric commonly used in ecology and macroevolution (Lum et al., 2022), and represents the amount of independent evolution (Felsenstein, 1985) contributed by a sequence to a tree.

**Figure 3.**
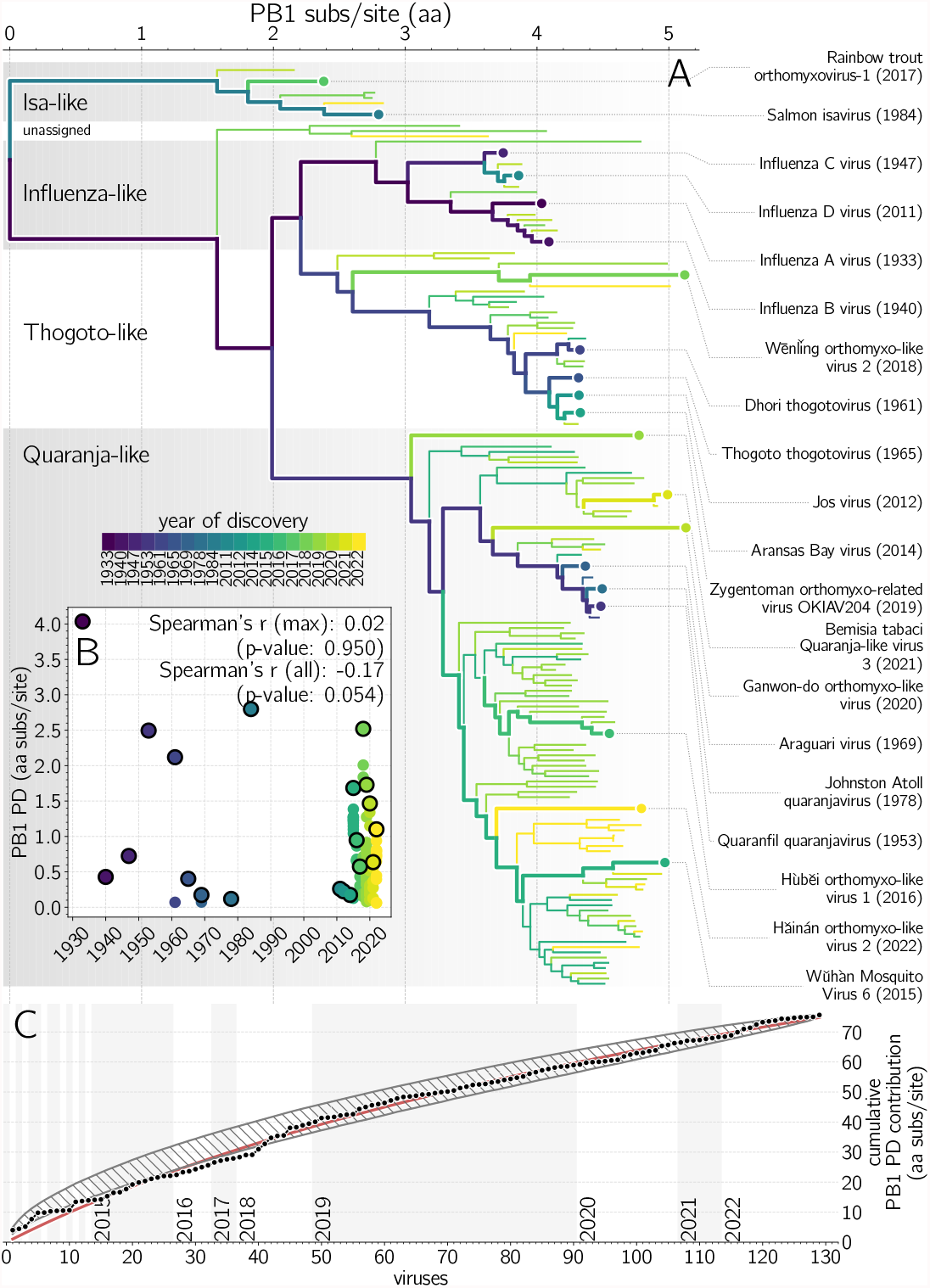
Discovery of orthomyxovirid PB1 phylogenetic diversity (PD). A) Maximum likelihood (ML) tree of orthomyxovirid PB1 proteins. Branch color indicates earliest discovery year of the lineage. Evolutionary history in purple branches was discovered earlier than yellow branches. Viruses contributing the most PD in their year of discovery are labelled on the right, and indicated with thicker paths in the phylogeny. B) Inset scatter plot shows PD contributions of each virus versus its year of discovery, with black outlines indicating the maximal PD contributor each year. While the average PD contribution of a newly discovered orthomyxovirus is decreasing with time (Spearman’s r=-0.17, one-tailed p-value: 0.027), the PD contribution of the most novel virus discovered each year has held steady (Spearman’s r=0.02, one-tailed p-value: 0.950/2=0.425). C) Cumulative PB1 PD contribution from successive orthomyxovirid discoveries (black dots) with logarithmic least-squares fit (red line). Gray hatched area indicates the 95 percentile range of cumulative PD contributions under 10000 random permutations of taxa discovery order.

We find that distinctive viruses, those contributing significant PD, have continued to be discovered each year (Fig. 3B). For example, the Wēnlǐng orthomyxo-like virus 2 found in 2018 is nearly as distinctive relative to the viruses discovered before it as the Infectious salmon anemia virus found in 1984 was. There is no correlation between the year of discovery and the maximum PD contributed by an orthomyxovirid (Spearman r = 0.02, p-value = 0.95). In contrast, the average PD contributed per virus does decrease with time (Spearman r = *−*0.17, one-tailed p-value = 0.054*/*2 = 0.027), as shared evolutionary history is attributed to earlier discoveries. While the orthomyxoviruses discovered each year are, on average, less distinctive, the increased host breadth and rapid pace of current studies result in evolutionarily highly distinctive viruses.

#### Projecting future PB1 phylogenetic diversity discovery

Fig. 3C shows the cumulative PD of PB1 after each new orthomyxovirid discovery. Early orthomyxovirid discovery efforts do show some bias, finding viruses more related to one-another than by chance: until 2018, the empirical accumulation of PD (black dots) is mostly below the 95 percentile envelope of 10000 random permutations of discovery order (gray hatched area). We fit the empirical data with a logarithmic function (*f* (*x*) = *A × log*_2_(1 + *x/B*), where A = 46.3 and B=62.4), indicated with a red line in Fig. 3C). We can extrapolate this curve into the future, *e*.*g*. 200th orthomyxovirid is expected to contribute *≈*0.26 amino acid substitutions per site to PB1 PD (bringing total PD to 95.9), and the 500th *≈*0.12 (total PD 146.8). Note that any remaining bias in the current viral discovery paradigm, leaving some parts of the *Orthomyxoviridae* tree of life undersampled, would manifest in a future PD curve higher than the extrapolation of the one observed so far. There is, regardless, an eventual limit, in which the PD gain of a new discovery is less than the threshold of difference used to define a new taxon (*e*.*g*. 0.1 aa sub/site would currently be reached around the 600th taxon). If current trends continue, there would remain at least hundreds of additional taxa to be discovered.

## 3 Discussion

### 3.1 Summary

In this work, we endeavored to show how the accumulation of metagenomic data can lead to a new stage in viral studies. We focused on the family *Orthomyxoviridae*, synthesizing data across numerous studies to analyze geographic, evolutionary, and diversity discovery trends.

We first focus on a single recently discovered mosquito RNA virus - Wǔhàn mosquito virus 6 (Li et al., 2015) (WuMV-6) - whose frequency and fast evolution uniquely enable the tracking of mosquito populations. In our experience, metagenomically discovered RNA viruses can be rare or, when encountered often, do not always contain sufficient signal to calibrate molecular clocks (Webster et al., 2015). WuMV-6 has rapidly disseminated across vast distances, and while anthropogenic (shipping, air travel) (Lounibos, 2002; Fonseca et al., 2006; Bataille et al., 2009) or abiotic (windborne migration) (Huestis et al., 2019) mechanisms may contribute, the virus’ extremely diverged and actively diversifying gp64 surface proteins suggest a potential vertebrate host. Indeed, the rapid sweep of the USA by West Nile virus was accelerated by the movement of both its mosquito vector and its diverse avian hosts (Di Giallonardo et al., 2015). While alternation between vertebrate host species could theoretically produce diversifying selection on the WuMV-6 surface protein, gp64 uses NPC1 (Li et al., 2019), a highly conserved metazoan protein, as its receptor. We thus believe it more likely that WuMV-6 gp64 diversity is selected for by repeat exposure to vertebrate hosts (Jong et al., 2007), which help disperse WuMV-6 (Lycett et al., 2019) and reduce its effective population sizes (Bedford et al., 2011). Previously, this sort of phylodynamic analysis was limited to known human and animal pathogens (Drummond et al., 2003; Wheeler et al., 2010). As metagenomic discovery efforts continue more systems like WuMV-6 will undoubtedly be found, contributing to research areas outside of virus evolution, like disease vector dispersal and shifting host distributions under climate change.

### 3.2 Molecular clock analyses applied to arthropod RNA viruses

Arthropod RNA virus population dynamics in the wild are not well understood at this time and, as such, the application of sophisticated phylogenetic analyses to these systems are only getting started. For example, *Drosophila* research to date has uncovered the full spectrum of dynamics - a suspected sweep of a Sigma virus in British *D. obscura* over roughly a decade (Longdon et al., 2011), the P element invasion of *D. melanogaster* that took place over a few decades (Anxolabéhère et al., 1988), and viruses of *D. melanogaster* and *D. simulans* whose common ancestry is on the order of hundreds of years (Webster et al., 2015). Similarly, vertical transmission (described in at least one member of the eight-segmented Asto-Ū sinis clade (Coatsworth et al., 2022)) and diapausing could also complicate molecular clock analyses of arthropod RNA viruses by affecting viral population dynamics and/or replication rates. As such, the identification of more arthropod RNA virus systems that are amenable to molecular clock analyses will be hugely important in illuminating the timescales of RNA virus population turnover in arthropods, identifying departures from neutral evolution, and providing much-needed context.

### 3.3 Long-term evolution of viral receptor binding and membrane fusion proteins

Zooming out, we find a highly modular and fluid genomic organization across members of the family *Orthomyxoviridae*. This presents an interesting conundrum – how and why are novel segments acquired so readily by orthomyxovirids, given that packaging signals encompass multiple sites (Baker et al., 2014) and obtaining segments via recombination is hard (Chare et al., 2003)? (Splitting genomes via segmentation is better documented (Kondo et al., 2006; Qin et al., 2014) and easier to explain conceptually (Ke et al., 2013).) Observed frequent switches in surface proteins may be selected for because of their importance in determining host and tissue tropism. We may also be missing additional classes of surface protein, beyond HEF-like and gp64, because of a reliance on sequence homology for protein identification. Indeed, many pooled studies produce incomplete viral genomes, with too few segments relative to their clade, so there is a possibility that significant undetected gain and loss of segments has occurred even within already-discovered viruses. To assess such evolutionary questions will require metagenomic studies to look beyond discovering conserved genes in increasing numbers of host species to completing viral genomes by sequencing individuals across geographic transects (Batson et al., 2021). Laboratory studies will be necessary to identify the functions of these novel segments and to confirm/determine tropism of discovered surface proteins (Arunkumar et al., 2021).

### 3.4 Predicting discovery of *Orthomyxoviridae* diversity

Finally, we assess the overall progress of orthomyxovirus discovery from the perspective of phylogenetic diversity (PD). We find that the many new orthomyxovirids being discovered every year are adding significant evolutionary history to their family tree. We may contrast this to the situation for birds, which have been studied and characterized for centuries and for which the discovery of new and distinctive species is now rare (Lum et al., 2022). Where the PD contribution of the most distinctive avian species discovered in each year exhibits a strong downwards trend (Fig. 4 from (Lum et al., 2022)), the PD of the most distinctive orthomyxovirus discovered each year remains high (our Fig. 3B). The aggregate trend also indicates that significant PD remains to be discovered: if logarithmic trend continues, known *Orthomyxoviridae* member diversity would double on the discovery of the 531st member. (While there is a risk that some metagenomic sequence represents endogenized viral genes, this is extremely rare for the RdRp gene we use to calculate PD (Whitfield et al., 2017).) This complements the argument made by (Parry et al., 2020) for the existence of many more influenza-like viruses based on virus-host codivergence and the existence of many unsampled host species. We believe that phylogenetic diversity measures, already in widespread use in ecology and macroevolution, will prove useful to the metagenomic virus discovery community as it seeks to assess ongoing progress and predict future payoff. One roadblock to the systematic application of these methods is the lack of consensus on how to define lineages as belonging to the same or different taxa. The International Commitee for the Taxonomy of Viruses (ICTV) has created a number of higher taxonomic degrees of organization (some of which are in dispute (Holmes and Duchêne, 2019)), while the growth of sequence databases has left many new viruses without an official taxonomic designation. Leadership in curating sequence databases can therefore have a disproportionately bigger impact on the direction the field of long-term virus evolution will take.

### 3.5 Keep going

This work was made possible by the public sharing of annotated genomes, raw sequencing data, and sampling metadata from groups across the world. As metagenomic surveys expand across diverse hosts and geographies, the accumulation of sequence data allows a depth of analysis that moves beyond virus discovery and into ecological and evolutionary dynamics; encountering new samples of previously seen viruses, instead of being seen as a disappointment, can be viewed as opportunity for more granular phylodynamic analysis. The evolutionary interdependence of sequence within and between organisms generates increasing returns on additional surveys. With appropriate study designs, good data organization, and public sharing strategies, the community’s search into the shape of the “virosphere” will offer large dividends for many fields of research.

## 4 Methods

### 4.1 Use of viruses for host tracking

Most of Wǔhàn mosquito virus 6 (WuMV-6) virus genome data (Chinese, Californian and Australian genomes) were derived from a previous publication (Batson et al., 2021). Assembled contigs from the Swedish study (Pettersson et al., 2019) were provided by John Pettersson while the Cambodian sequences were kindly provided by Jessica Manning, Jennifer Bohl, Dara Kong and Sreyngim Lay, where WuMV-6 segments described later (Batson et al., 2021) were identified by similarity.

Puerto Rican segments were recovered by mapping reads from SRA entries SRR3168916, SRR3168920, SRR3168922, and SRR3168925 (Frey et al., 2016) to segments of Californian strain CMS001 038 Ra S22 using bwa v0.7.17 (Li and Durbin, 2009) but most segments except for NP did not have good coverage to be assembled with certainty. Greek segments were recovered by mapping reads from SRA entry SRR13450231 (Konstantinidis et al., 2021) using the same approach as described earlier. New Chinese segments from 2018 (He et al., 2021) were similarly recovered by mapping reads from China National GeneBank Sequence Archive accessions CNR0266076 and CNR0266075 using the same approach as described earlier. New Chinese and Greek segments tended to have acceptable coverage except for segments hypothetical 2 and hypothetical 3 where only individual reads could be detected. The presence of WuMV-6 in more locations around the world was determined based on the presence of reads with *≥*90% identity to WuMV-6 PB1 amino acid sequence AJG39094 in Serratus (Edgar et al., 2022). In this way the presence of WuMV-6 was detected in Connecticut, Trinidad and Tobago, Tunisia and Philippines.

All successfully assembled segments (from China, Australia, Cambodia, California, Greece, and Sweden) were aligned using MAFFT (Katoh et al., 2005) and trimmed to the coding regions of each segment. PhyML v.3.3.2 was used to generate maximum likelihood phylogenies of each segment under an HKY+Γ_4_ (Hasegawa et al., 1985; Yang, 1994) model. Each tree was rooted via least squares regression of tip dates against divergence from root in TreeTime (Sagulenko et al., 2018).

To confirm sufficient molecular clock signal in WuMV-6, maximum likelihood phylogenies (Stamatakis, 2014) were inferred from all available aligned segments under a GTR+Γ model and rooted via least-squares regression using TreeTime (Sagulenko et al., 2018) and visualized using root-to-tip plots for each segment. The genome-wide root-to-tip plot was made by only focusing on strains for which we had complete genomes and summing the divergence of each sequence from the root of their segment tree for any given strain.

We also ran molecular clock analyses on all available segment sequences individually using BEAST v.1.10.4 (Suchard et al., 2018) with tip date calibration, GTR+Γ_4_ nucleotide substitution model, strict molecular clock, and a constant population size tree prior. The date of collection for strain QN-3 (the first WuMV-6 strain to be discovered) was sampled uniformly from the interval between years 2013 and 2014. Analyses were run for 50 million states, sampling every 5 000 states. With convergence confirmed visually in Tracer v.1.7.1 (Rambaut et al., 2018) and 10% of the states discarded as burn-in before summarizing the posterior distribution of trees using TreeAnnotator with the common ancestors option (Heled and Bouckaert, 2013). To confirm the diverged WuMV-6 sequences from Sweden were not influencing the molecular clock rate we repeated the entire analysis but excluded the samples from Sweden.

27 complete WuMV-6 genomes (13 from California, 7 from Australia, 3 from Cambodia, 3 from Sweden, and 1 from China) were analyzed using the reassortment network method (Müller et al., 2020) implemented in BEAST v2.6 (Bouckaert et al., 2019). For the smallest segment coding for the hypothetical 3 protein, two Ns were inserted after the 349th nucleotide from the initiation codon ATG to account for the presence of a suspected splicing site (Batson et al., 2021) that brings a substantial portion of this segment back to being coding. Each segment was partitioned into codon positions 1+2 and 3 evolving under independent HKY+Γ_4_ (Hasegawa et al., 1985) models of nucleotide substitution and independent strict molecular clocks calibrated by using tip dates. By default a constant effective population size coalescent tree prior is applied to the reassortment network. Default priors were left in all cases except for effective population size (set to exponential distribution with mean at 100 years) and reassortment rate (set to exponential distribution with mean 0.001 events/branch/year) to get conservative estimates and prevent exploration of complicated parameter space. MCMC was run for 200 million states, sampling every 20000 states in triplicate, after which all chains were combined after discarding 10% of the states as burn-in and confirmed to have reached stationarity using Tracer (Rambaut et al., 2018). The reassortment network was summarized using the native BEAST v2.6 tool (ReassortmentNetworkSummarizer) provided with the package. Posterior embeddings of each segment within the network (in the form of clonal phylogenetic trees) were summarized using TreeAnnotator v.1.10.4 after combining independent runs after discarding 10% of the states as burn-in.

In our personal experience ReassortmentNetworkSummarizer is overly conservative when summarizing reassortment networks due to reassortant edges requiring conditioning on both the origin and destination clades. As such we removed reassortant edges with ≤0.1 posterior support and summarized posterior supports by first extracting the embedding of each segment from the summarized network by carrying out the subtree prune-and-regraft procedures implied by reassortant edges and then finding how many of the same clades are found in posterior summaries of segment embeddings and how many of those are supported with posterior probability ≥0.95 to produce Figure 1. Similarly, the 95% highest posterior density interval for most recent common ancestor of the “Pacific rim” clade (*i*.*e*. genomes collected outside of Sweden) produced by ReassortmentNetworkSummarizer in Figure 1 do not overlap perfectly with estimates we extracted from the posterior distribution of reassortment networks though.

All trees were visualized using baltic (https://github.com/evogytis/baltic) and matplotlib (Hunter, 2007).

### 4.2 Orthomyxovirus segmentation and surface proteins

For each clonal WuMV-6 segment embedding within the reassortment network 1 000 trees from the posterior distribution were extracted after removing 10% burnin and combining all three independent runs. These trees were then used as empirical trees to be sampled from in a BEAST v.1.10.4 (Suchard et al., 2018) renaissance counting analysis (Lemey et al., 2012) run for 10 million states, sampling every 1 000 states.

Orthomyxovirid PB1 protein sequences from each genus - isa-, influenza, thogoto-, and quaranjaviruses were used as queries in a protein BLAST (Altschul et al., 1990) search with influenza A, B, C, and D viruses excluded from the search. Having identified the breadth of PB1 protein diversity and having downloaded representative PB1 proteins of influenza A, B, C, and D viruses we aligned all sequences using MAFFT (Katoh et al., 2005) (E-INS-i mode) and removed sequences that were identical or nearly identical, as well as short or poorly aligning sequences. We repeated this procedure with blast hits to capture as much PB1 diversity as is publicly available. Partial, poorly aligning or insufficiently distinct PB1 sequences were removed from the analysis.

We used the same data gathering technique for surface proteins. To identify HEF-like proteins we used isavirid HE and influenza C and D virus HEF proteins as queries but did not identify any additional proteins. The claimed hemagglutinin proteins of Rainbow and Steelhead trout isaviruses did not resemble anything on GenBank except each other and did not produce any significant hits via HHpred (Finn et al., 2011). BLAST searches using orthomyxovirid gp64 relatives identified thogoto- and quaranjavirus surface proteins, as well as baculoviruses and rhabdoviruses with identifiably related proteins. The presumed gp64 proteins found within the clade encompassed by Ū sinis, Astopletus (discovered in California) and WuMV-6 with Guadeloupe mosquito quaranja-like virus 1 (previously described), referred to here as the Asto-Ū sinis clade within quaranjavirids, did not resemble anything on GenBank via protein BLAST but were all inferred to strongly resemble gp64 proteins via HHpred and as such were aligned using MAFFT in G-INS-i mode.

The PB1 dataset was then reduced to viruses for which gp64 sequences were largely available, members of the Asto-Ū sinis clade, and more diverged members. Phylogenetic trees for both PB1 and gp64 proteins were inferred using PhyML v.3.3.2 and rooted on isavirids for PB1 sequences and depicted unrooted for gp64.

For each PB1 blast hit we searched GenBank for the rest of the genome, ignoring any genomes that appear to have fewer than 6 segments on account of the three RdRp segments, nucleoprotein and occasionally surface proteins being far easier to identify and all of the best-studied orthomyxovirids having at least six segments. We visualized PB1 and gp64 trees using baltic and annotated tips with number of segments identified and category of surface protein used, where available. For annotating genome organization we further marked the earliest plausible common ancestors that must have possessed a given genome organization and highlighted all of their descendants as a prediction for which other datasets might have the missing segments.

### 4.3 Phylogenetic diversity estimation

The larger PB1 sequence data set (prior to reduction) was used to infer a maximum likelihood tree using PhyML v.3.3.2 which was rooted on isavirids. For each protein, the date of either its publication in literature or on GenBank was noted. For each year of discovery available, tree branches were marked with the evolutionary path uncovered that year, starting from oldest published sequences. The sum of branch lengths contributed by any given sequence to the tree is what we call phylogenetic diversity (PD). As well as the relationship between year of discovery and maximum PD contributed in Fig. 3A we looked at successive and unique PD contributions by each newly discovered orthomyxovirid in comparison to a neutral PD discovery curve.

### 4.4 Data availability

Data and scripts to replicate analyses are publicly available at https://github.com/evogytis/orthomyxo-metagenomics.

## 5 Acknowledgements

GD acknowledges the support of European Molecular Biology Organization (EMBO) installation grant IG-5305-2023. We would like to acknowledge the contributions of Amy Kistler, Maira Phelps, Cristina Tato, and Fabiano Oliveira in setting up sample logistics, experimental design, and data analysis for the Californian mosquito virome study. We would like to thank Darren Obbard for numerous and fruitful discussions. We are grateful to Jessica Manning, Dara Kong, Sreyngim Lay, Alex Greninger, Mang Shi, Eddie C Holmes, Dana Price, and John Pettersson for sharing assembled sequence data.

## Notes

https://github.com/evogytis/orthomyxo-metagenomics

### Competing Interest Statement

The authors have declared no competing interest.

### Summary of Updates

Additional supplementary figures were added to show WuMV-6 genomes contain sufficient molecular clock signal on their own, some clarification added to the text.

