## Supplemental materials for "Accumulated metagenomic studies reveal recent migration, whole genome evolution, and undiscovered diversity of orthomyxoviruses"

### 1 Supplementary material

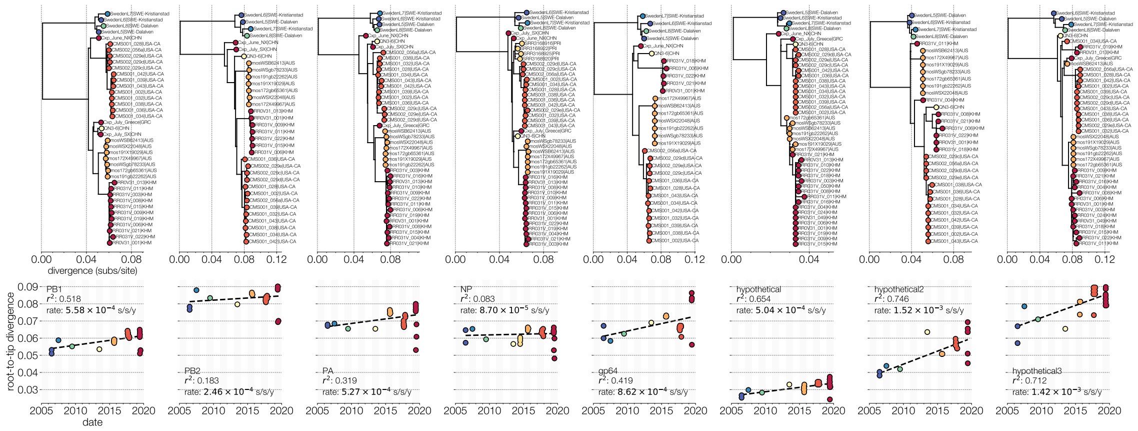


**Figure S1. Root-to-tip plots for all segments**. Top row shows the maximum likelihood phylogenetic trees of WuMV-6 segments rooted via least-squares regression. Bottom row shows the root-to-tip plots, the correlation between collection date (x-axis) and divergence of each tip from the root of the tree (y-axis). A dotted line indicates the root-to-tip regression line with slope (*i.e.* evolutionary rate esimate) and the $r^2$ value displayed either in the upper left or the bottom left corner of each plot. Points are colored by the collection date of each sample.

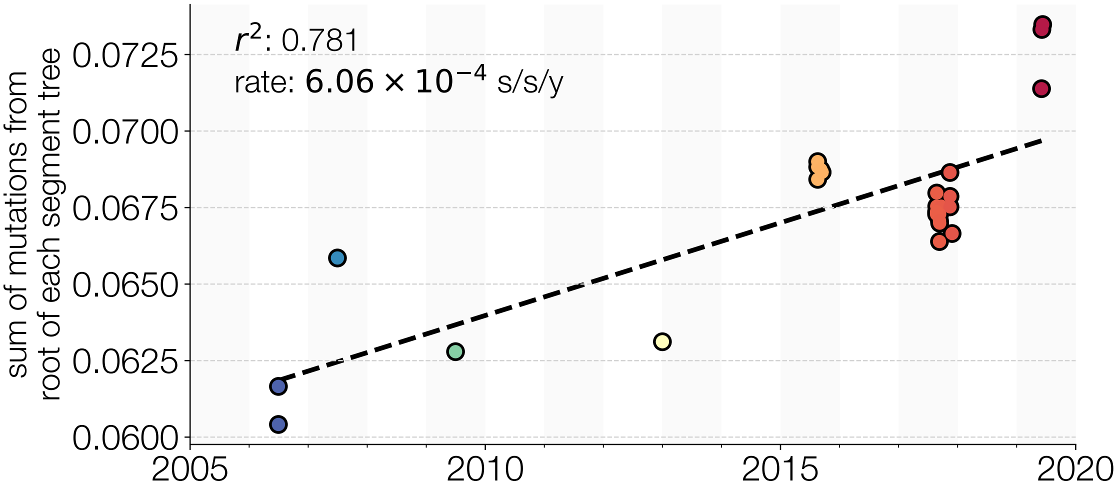


**Figure S2. Root-to-tip plot for whole genomes**. For each tip in segment trees rooted via least-squares regression in Supp. Fig. S1 (if the entire genome was available) we first converted their height from subsitutions per site to numbers of mutations (by multiplying tip height by segment length). We then summed these mutations across segments to arrive at a genome-wide distance of each tip from the root. We finally divided this genome-wide distance by total WuMV-6 genome length to arrive at a genome-wide divergence of each tip expressed in subsitutions per site. A dotted line indicates the root-to-tip regression line with slope (*i.e.* evolutionary rate esimate) and the $r^2$ value displayed in the upper left corner. Points are colored by the collection date of each sample.

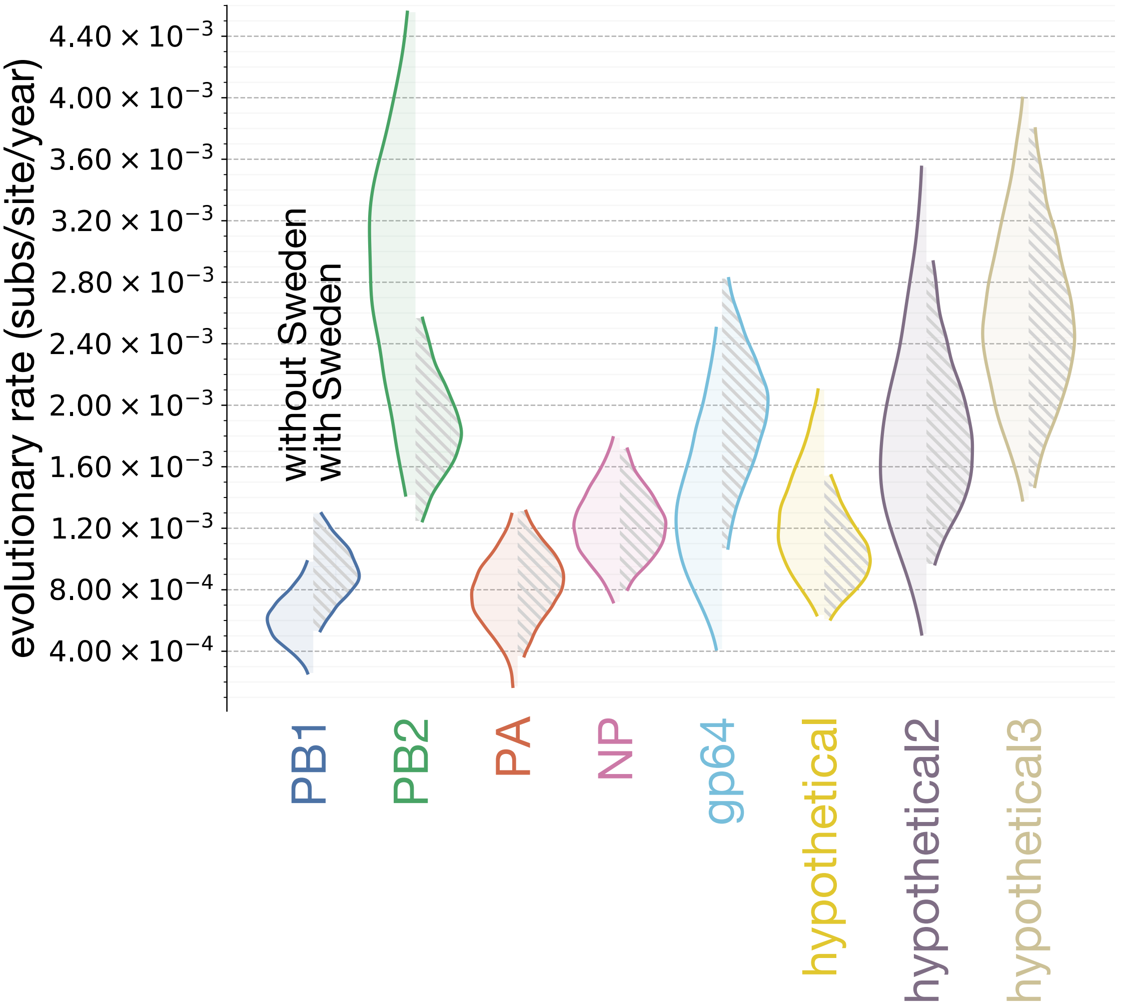


**Figure S3. Evolutionary rate estimates from tip-calibrated molecular clock analyses**. Half-violin plots indicate the 95% highest posterior density (HPD) intervals for evolutionary rate when segment sequences shown in Supp. Fig. S6 are used for molecular clock calibration individually with an uninformative prior on the molecular clock rate. The density on the right shows the evolutionary rate estimate when the Swedish samples that comprise an outgroup are excluded while the hatched density on the left shows the estimate when all available sequences are included. Densities are colored according to segment.

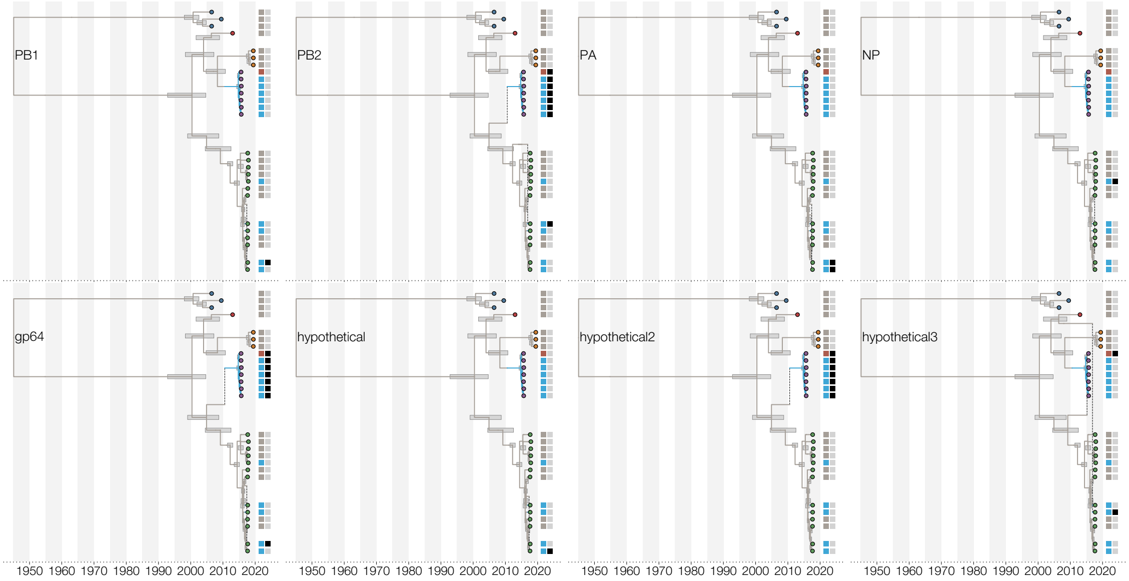


**Figure S4. Segment paths through the reassortment network**. Each panel shows the embedding of every segment in the reassortment network of Wǔhàn mosquito virus 6 genomes. Tips are colored by location, as in Figure 1A in the main text. The first colored box to the right of the tree indicates how many successive reassortments the entire genome a tip belongs to has experienced reassortment - cycling between grey, blue, and brown colors for each successive event. The second colored box indicates whether the segment in question has undergone a reassortment event. Horizontal gray bars indicate node height 95% HPD intervals in the summary network.

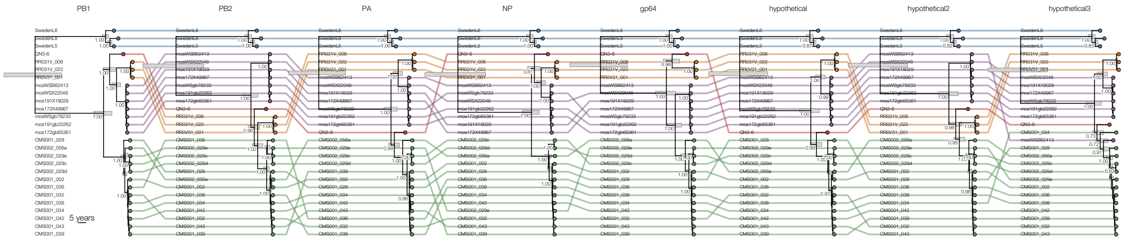


**Figure S5. Tangled chain of maximum clade credibility trees of individual segment embeddings**. Each tree corresponds to a segment of Wǔhàn mosquito virus 6, with tips colored the same as Fig. 1 in the main text. The same tips are connected with colored lines between successive trees to indicate changes in their phylogenetic position, their names are given at the base of each tree. Numbers at each internal branch longer than 1.0 year correspond to the node's posterior probability. Horizontal gray bars indicate node height 95% HPD intervals.

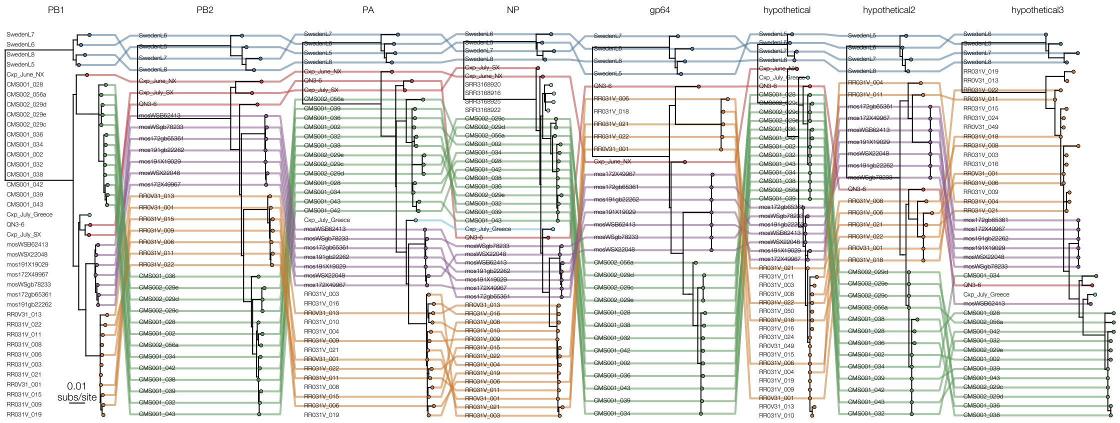


**Figure S6. Tangled chain of maximum likelihood trees of individual segments**. Each tree corresponds to all available segments of Wǔhàn mosquito virus 6, with tips colored the same as Figure 1A in the main text. The same tips are connected with colored lines between successive trees (if the corresponding segment was identified for that genome) to indicate changes in their phylogenetic position, their names are given at the base of each tree. Unlike Supp. Fig. 2, samples from Puerto Rico, Greece, Cambodia, and additional Chinese sequences are included here.

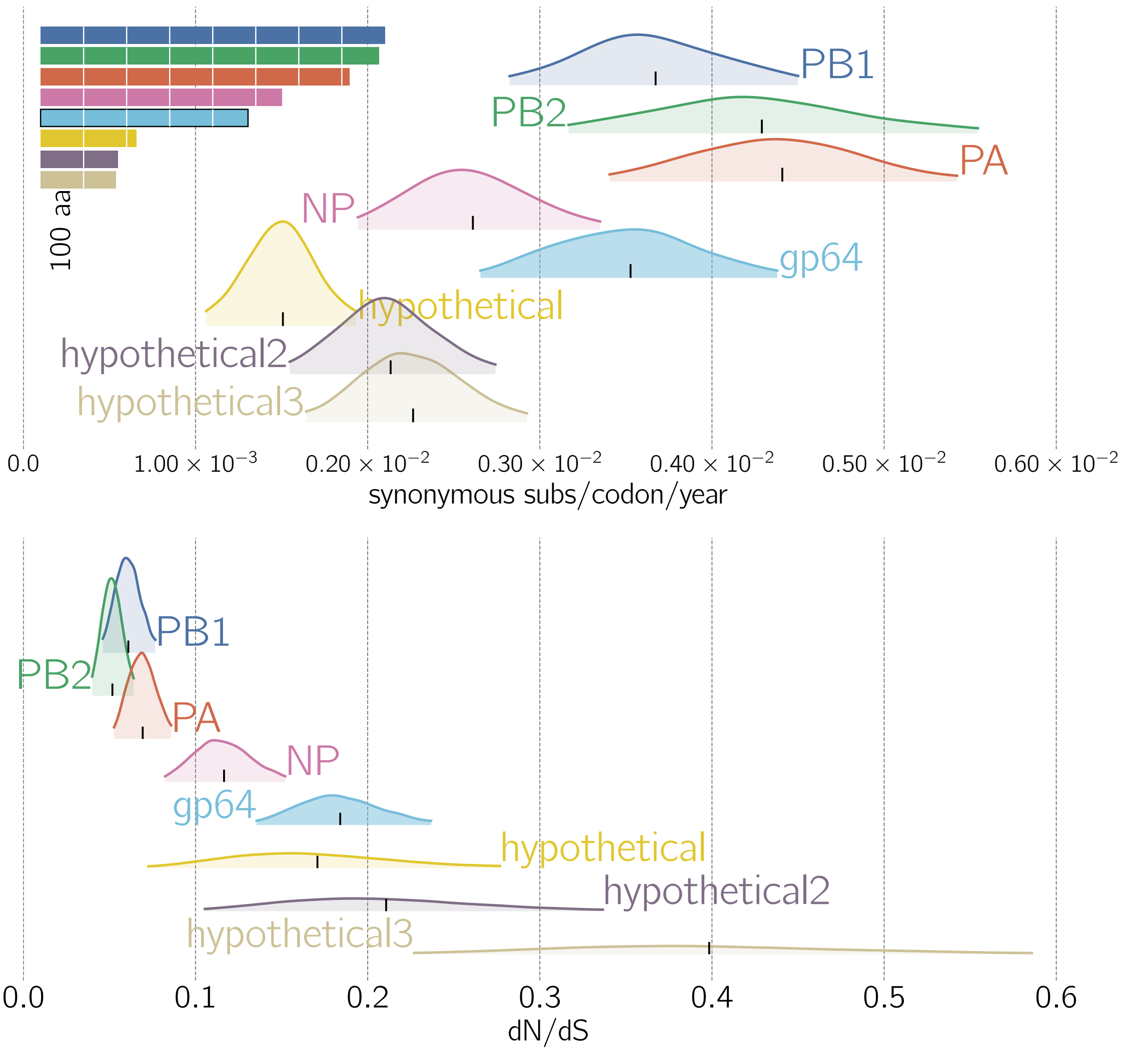


**Figure S7. Estimates of synonymous rate of evolution and dN/dS for Wŭhàn mosquito virus 6 genes**. The top panel shows 95% highest posterior densities of the synonymous rate of evolution for each WuMV-6 gene (distinguished by color). Colored rectangles in the top left show the relative lengths of each gene with white lines indicating 100 amino acid increments. The bottom panel shows 95% highest posterior densities of dN/dS for each WuMV-6 gene.

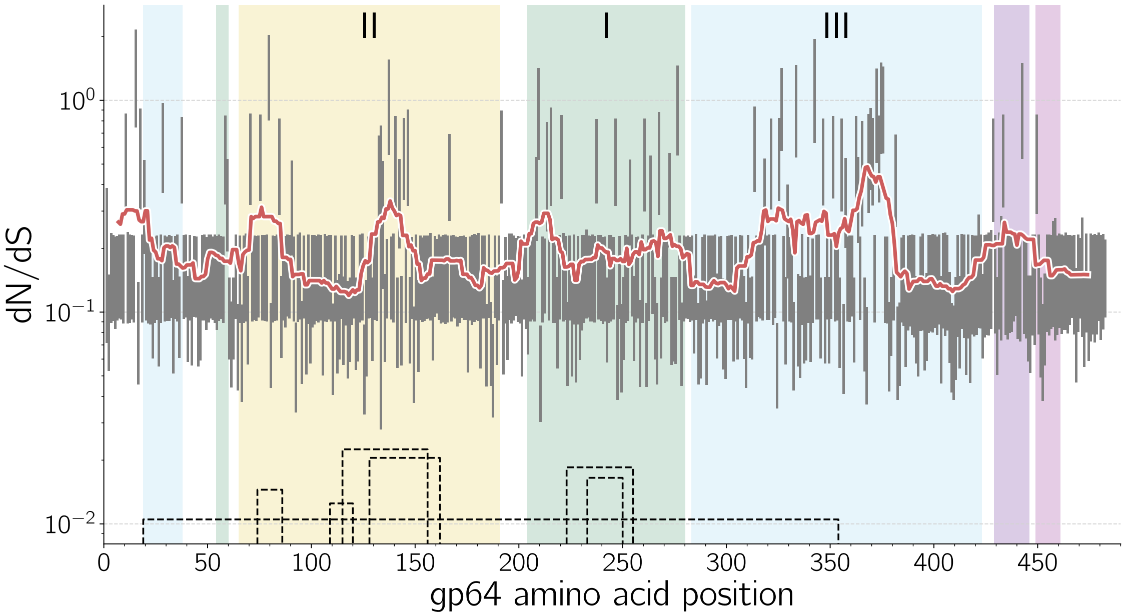


**Figure S8. Estimates of dN/dS in gp64 per codon**. Grey vertical lines indicate the 95% highest posterior density interval for dN/dS per codon in gp64 of Wǔhàn mosquito virus 6. The red line tracks mean estimated dN/dS in a sliding window of 15 amino acids. Regions of the protein are colored and labelled according to nomenclature proposed by (Garry and Garry, 2008). Dashed lines at the bottom depict putative disulfide bonds between cysteins in gp64, based on (Garry and Garry, 2008). The two peaks in average dN/dS in domain II occupy predicted regions of thogotovirus gp64 fusion loops which are inserted into the target membrane. A notable region of elevated dN/dS is seen in domain III but is difficult to explain since the region is expected to be proximal to the viral, and not the host, membrane.

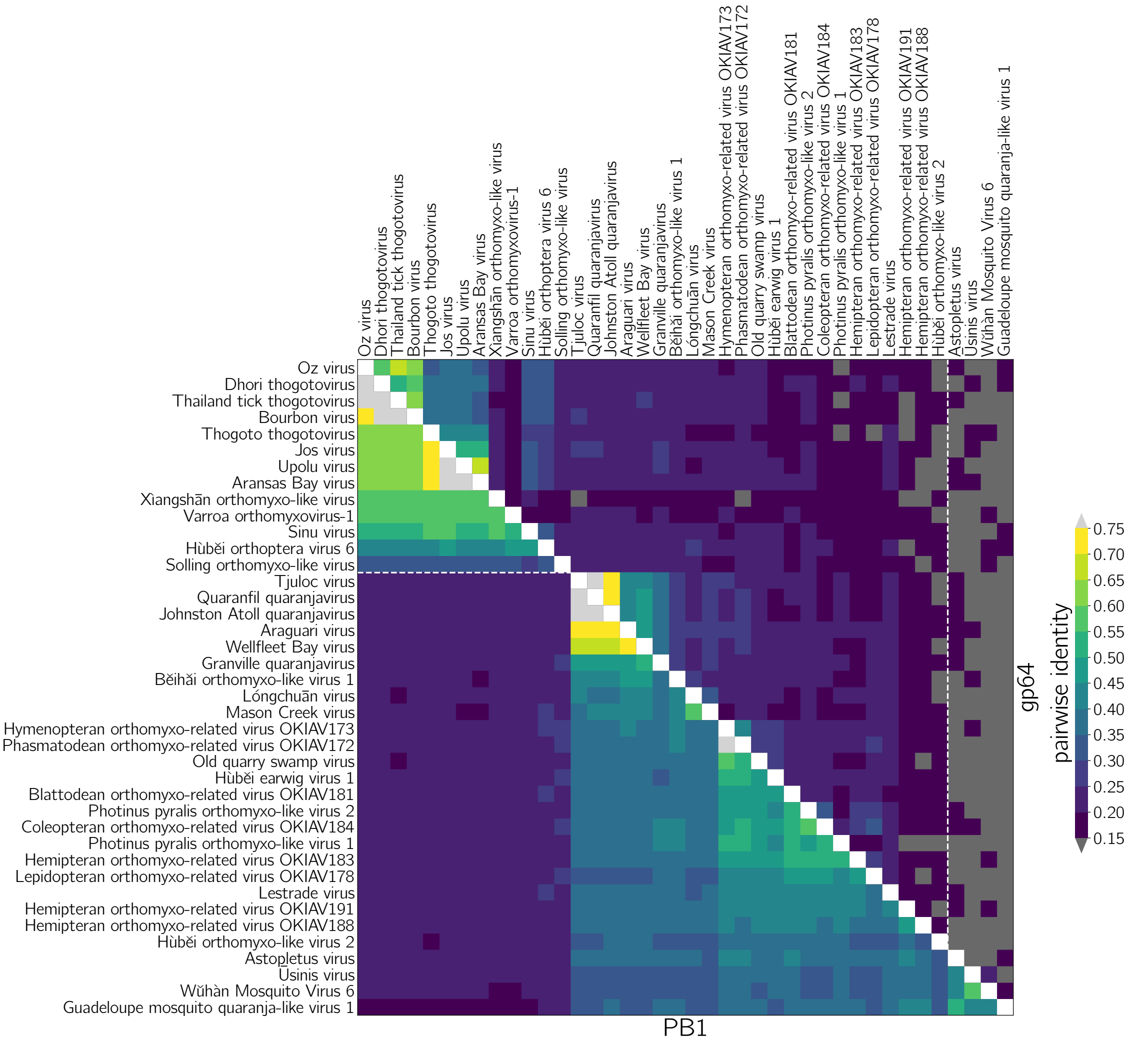


**Figure S9. Pairwise distance matrix between PB1 (lower triangle) and gp64 (upper triangle) proteins**. Dashed horizontal line marks the difference in PB1 between thogoto-like (above) and quaranja-like (below) viruses. Dashed vertical line marks the difference between stereotypically *Quaranjavirus* (to the left) and the markedly diverged quaranja-like gp64 proteins used by the eight-segmented Asto-Ūsinis clade (to the right).

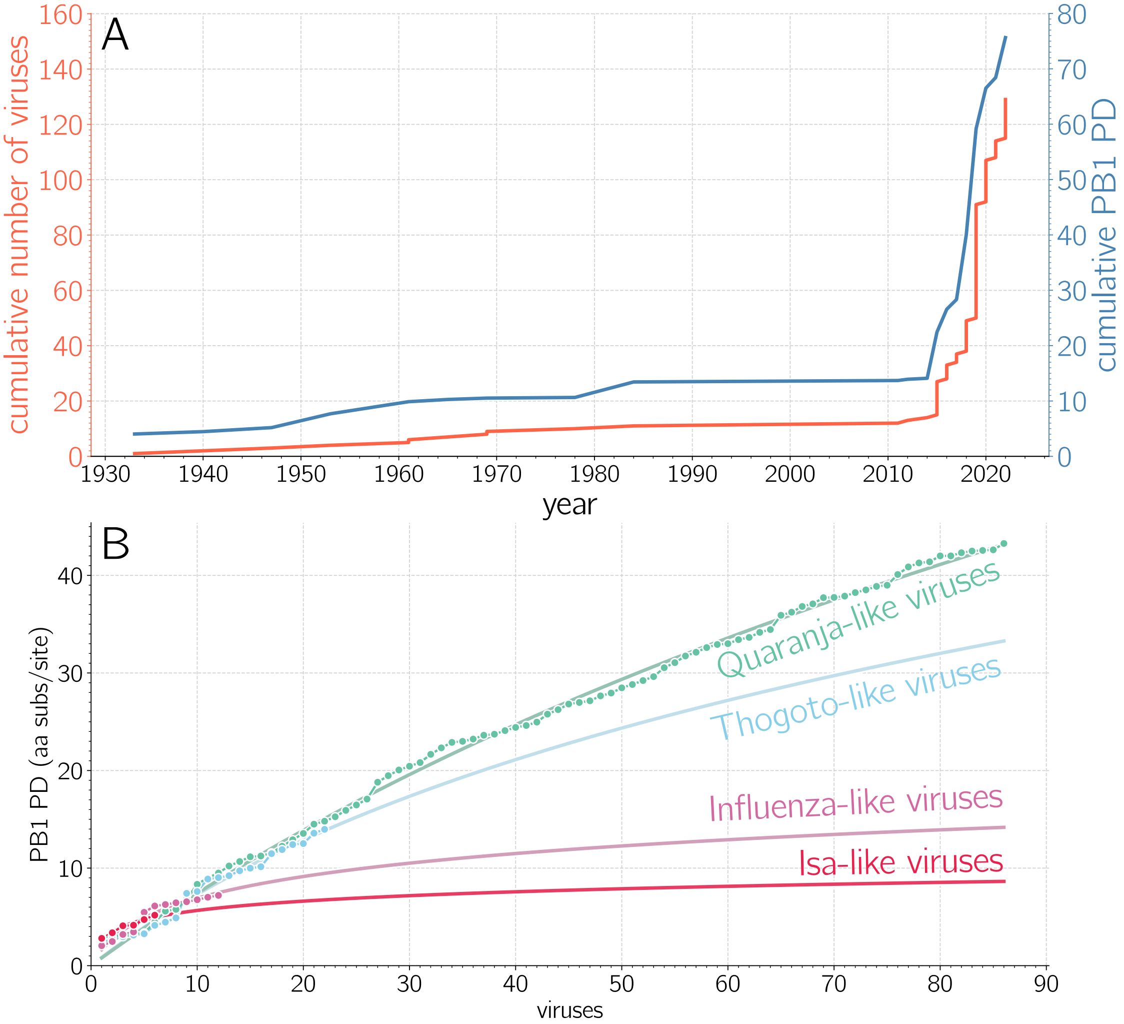


**Figure S10. Orthomyxovirus PB1 phylogenetic diversity discovery**. A) Orthomyxovirus (red) and PB1 phylogenetic diversity (blue) discovery curves over time. Regardless of whether discovery efforts are quantified in terms of distinct virus numbers or phylogenetic diversity, 2015 has marked a major turning point for discovery of orthomyxo- and other RNA viruses. B) Cumulative orthomyxovirus PB1 diversity discovery at the level of clades. Colored points track the cumulative diversity discovery (*i.e.* same as Fig. 3C) across well-defined orthomyxovirus clades (Quaranja-, Thogoto-, Influenza-, and Isa-like viruses) extracted from the PB1 tree shown in Fig. 3A. Logarithmic saturation curves are extrapolated for each clade and annotated with their name on the right.
